# Comparison of classification accuracy and feature selection between sparse and non-sparse modeling of metabolomics data

**DOI:** 10.1101/2023.04.03.535336

**Authors:** Arisa Toda, Misa Goudo, Masahiro Sugimoto, Satoru Hiwa, Tomoyuki Hiroyasu

## Abstract

Machine learnings such as multivariate analyses and clustering have been frequently used for metabolomics data analyses. In metabolomics data analyses, how much difference there is between the results calculated by supervised and unsupervised learning models is an interesting topic. Since metabolomics data include hundreds to thousands of metabolites greater than the sample numbers, only a small fraction of metabolites is relevant to the phenotype of interest. For this reason, sparse mechanisms have been introduced into many machine learning models. However, its explanatory power decreases when the number of explanatory variables is reduced to an extreme level. In this paper, serum lipidomic data of breast cancer patients (1) pre/post-menopause and (2) before/after neoadjuvant chemotherapy was chosen as one of metabolomics data. Here, this data was analyzed by partial least squares (PLS) for regression and K-means and hierarchical clustering for clustering. Results were also compare with the sparse modeling. Between the non-sparse and sparse modeling accuracy, there is no significant difference. Metabolite subsets selected by sparse modeling were almost identical to the PLS-selected features. At the same time, several metabolites were consistently selected regardless of the algorithm used. These results contribute to exploring biomarkers in high-dimensional metabolomics datasets.

## 1 Introduction

Recent advances in *omics* technologies have allowed us to fully quantify molecules in living organisms and generate extensive data, such as proteomes and transcriptomes [30]. Observing changes in hundreds to thousands of molecular-level networks has enabled a comprehensive understanding of the etiology and pathogenesis of various diseases [49], exploration of diagnostic and prognostic markers, and the search for therapeutic targets based on patterns of multiple molecules rather than single molecules [35].

Metabolomics is a comprehensive and simultaneous molecular profiling method that captures holistic metabolic pathways. Nuclear magnetic resonance and mass spectrometry (MS) are widely used in metabolomics. These techniques are increasingly sensitive, enabling the simultaneous measurement of hundreds to thousands of metabolites. Furthermore, the highly sensitive MS has enabled the quantification of a wide variety of metabolites, which has increased the dimension of observed information [27, 40].

Machine learning algorithms include regression (as a supervised method) and clustering (as an unsupervised method) [1, 5]. Both of these algorithms are used to interpret multidimensional *omics* data. Supervised methods are used to identify metabolite patterns according to known phenotypic labels. Partial Least Squares (PLS) regression [57], one of the most used algorithms in metabolomic data analyses, is a projection method in which multivariate data are projected into a latent space and then regressed to dependent features [17]. This method effectively analyzes metabolomics data with high covariance [13]. Support Vector Machine (SVM) [2] and Random Forest (RF) [4] are also commonly employed. In particular, RF has been reported to be effective for overfitting problems in high-dimensional metabolomics data [6]. On the other hand, unsupervised methods use unlabeled data to identify structures and patterns in *omics* data. Hierarchical clustering (HC) [28] and K-means clustering [23] are the most widely used unsupervised methods [12, 33, 39]. Although clustering is unsupervised, the clusters generated can be used as features in supervised machine learning algorithms.

The high throughput of metabolomics data poses a statistical problem wherein the number of experimental units *n* is small, but the feature dimension p is large [29]. Moreover, metabolomics data also have the problem of multicollinearity, in which features are highly correlated. Therefore, increasing the number of features without consideration is not advisable. One approach to solve this problem is sparse modeling, which imposes sparsity simultaneously with training. This has the potential to identify the minimum set of most informative features (i.e., the least optimal problem). However, to our knowledge, very little is known about the effectiveness of sparse modeling compared to the classification performance and feature selection techniques of traditional benchmark algorithms in metabolomic studies. In a previous study, five unsupervised learning algorithms were compared using metabolomics data, including sparse modeling [18]. However, the comparisons were limited to the clustering algorithm, and performance evaluation was limited to internal validation based on silhouette values.

One of the final goals of metabolomics research is to identify biomarkers that strongly represent phenotypes. Therefore, in this context, the interest is not in the classification of the samples but in the features. Many feature selection methods for biomarker identification have been proposed and discussed in various literature [21, 45, 48]. In some algorithms, such as Variable Importance in Projection (VIP) in PLS and Gini coefficients in RF, corresponding model metrics are used to rank features after training. In contrast, K-means and HC lack such metrics. Sparse model clustering (e.g., sparse K-means clustering [55]) allows the interpretation of metabolites that contribute to clustering in a subset of features. Partitioning Around Medoids (PAM) is a similar technique that introduces a dissimilarity matrix. Single-layer neural network methods, such as Self-Organizing Maps (SOM) [34], have also been developed. These algorithms classify the given samples according to their similarity, rank the importance of observed features based on their clustering contribution, and use the top-ranked features for biochemical considerations and markers. However, all these algorithms still use all observed features.

The higher the dimensionality and complexity of the data, the lower the reliability and interpretability of feature importance obtained a posteriori from the learning algorithm. In order to address this problem, it is helpful to remove phenotypically irrelevant features from the analysis. Under minimum feature subsets, it is possible to avoid overfitting scenarios in training and yield a learner with better generalization performance. Further, this reduces the computational cost and improves the interpretability of the outputs. Feature selection methods can be categorized into three approaches: filter, wrappers, and embedded [22, 60]. The filter method (i.e., removing low-quality features before training the algorithm according to feature ranking) is the most straightforward and widely used feature selection approach in metabolomics research. However, two concerns are noted with this method. First, although the subset size of potential features is essential, the filtering method is subjective because it requires selecting a cutoff value and manually determining the strength of the filter. Second, as feature selection is performed before the build of the learner, the selected features may not be optimal for improving the performance of the learner. In contrast, embedding methods such as sparse modeling, which implement the learning and feature selection steps simultaneously and may be useful as a feature selection tool for biomarker discovery. However, such methods have not been fully exploited in metabolomic research. Although embedding methods execute feature selection based on a single learning algorithm, some studies have reported its excellent performance with different models [11, 50].

This study aims to compare sparse modeling among conventional benchmark algorithms. Specifically, we aim to comprehensively compare sparse K-means [55] and sparse PLS regression [8] with five algorithms conventionally used in metabolomics in terms of prediction performance, computational cost, and selection of features that highly contribute to discrimination. Clustering and regression were applied to the binary classification problems in this study. For the two sparse modeling methods, trends in the variability of feature subset sizes selected were investigated using multiple trials. Finally, the quality of the feature subset selected by sparse modeling was compared with that of the filter method. Two numerical experiments using serum lipidomic data from breast cancer patients showed that sparse modeling achieved classification accuracy comparable to or better than non-sparse modeling, despite the use of feature subsets. Furthermore, most feature subsets selected by sparse modeling overlapped with the feature selection based on VIP using the commonly used PLS. The selected metabolite subsets showed classification accuracy comparable to that of feature filtering.

## 2 Materials and Methods

We compared three clustering and four regression algorithms, including sparse modeling, in terms of classification performance, computational cost, metabolites contributing to discrimination, and feature selection. The selected clustering and regression algorithms were well-known basic algorithms. A benchmark metabolite dataset (MTBLS92 [24]) was used for the numerical experiments. Each algorithm is briefly described in the following, followed by an overview of the dataset, preprocessing, and analysis.

### 2.1 Mini-review of clustering and regression algorithms frequently used in metabolomics research

This section reviews a brief overview of the available algorithms. Figure 1 shows a diagram of these algorithms.

**Figure 1.**
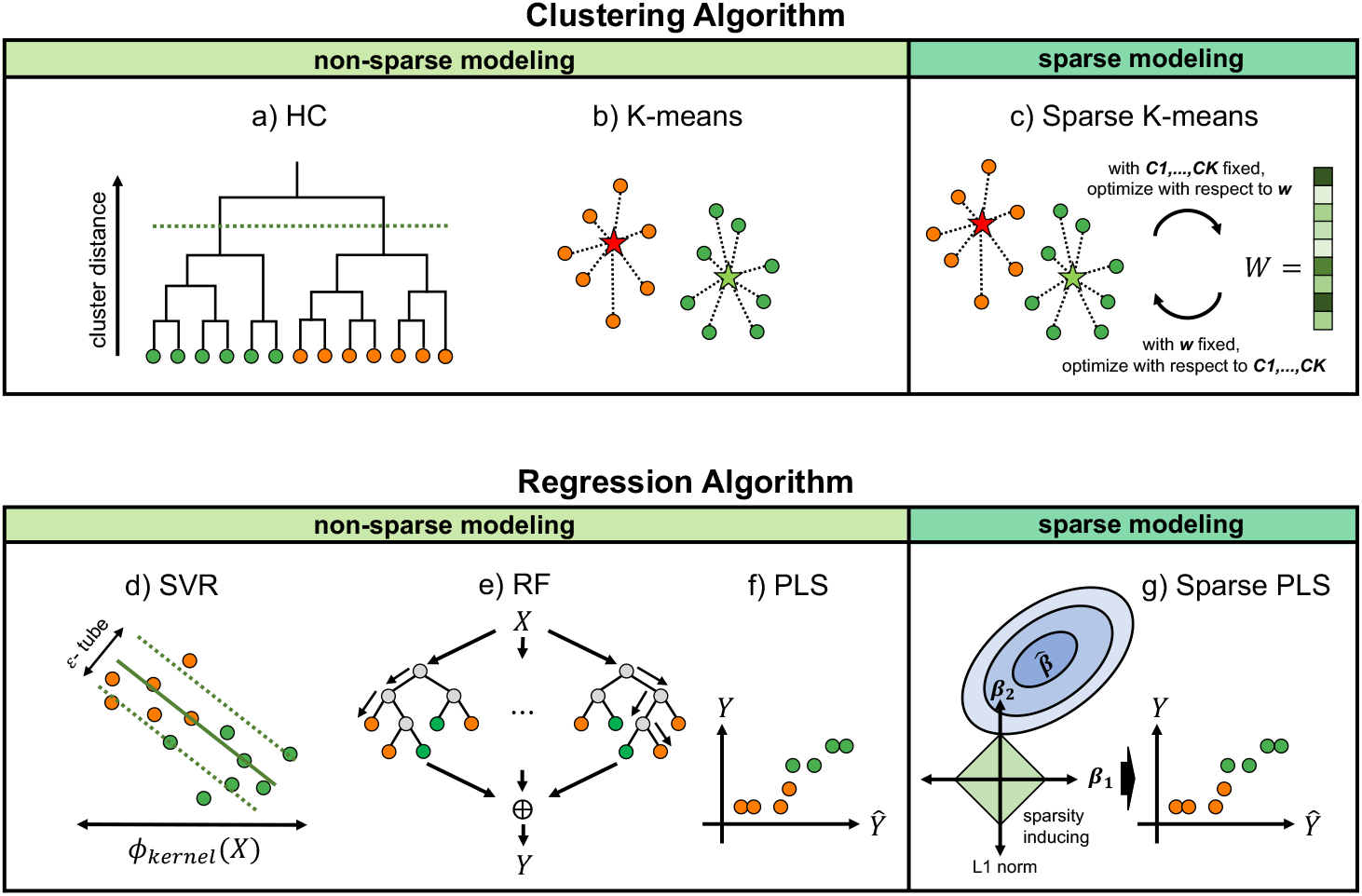
Schematic illustration of the compared algorithms. Orange and green dots represent sample data with the class labels. (a) Hierarchical clustering (HC): a clustering algorithm that finds the most similar (or dissimilar) combinations of sample data and groups them in order. (b) K-means: a non-hierarchical clustering algorithm. The average of the clusters is used to divide the data into predefined k clusters. (c) Sparse K-means: a sparse modeling version of K-means clustering, where iterative optimization is performed on a subset of useful features. (d) Support vector regression (SVR): a pattern identification algorithm for binary classes. The error and weight coefficients are minimized after transforming the data space using liner or nonliner kernel function. (e) Random forest (RF): an ensemble learning algorithm that constructs many decision trees by random sampling with overlap and takes the average of the predictions for each tree. (f) Partial least squares (PLS): a discrimination algorithm that uses principal component regression to extract principal components from the data to maximize the covariance between the explanatory and objective features. (g) Sparse PLS: sparse modeling with PLS that reduces the discriminant features by imposing sparsity on direction vectors via a *L*_1_ penalty.

Three clustering algorithms were used: a) HC, b) K-means, and c) sparse K-means. a) HC is used in many studies because it does not require a predefined number of clusters, unlike K-means clustering, and because it is possible to graphically represent the clustering results as a tree that summarizes the proximity and classification relationships within a set of data. Implementation requires the pre-definition of a linkage function (such as single linkage, average linkage, full linkage, or Ward linkage) that defines the distance between any two clusters. There are two approaches to HC: the top-down approach, called the agglomerative method, and the bottom-up approach, called the divisive method. In practice, the aggregative method is used more widely. It is a method that first considers all data points as clusters and repeats the merging of neighboring clusters according to a predefined distance until there is only one cluster.

b) K-means clustering first assigns all sample data to random clusters and then searches for the best partition based on the distance from the cluster center of gravity. The number of clusters, *k,* must be specified in advance. c) Sparse K-means clustering alternates between clustering and feature weighting optimization, where each feature is assigned a weight of ≥ 0. Sparse K-means calculates the sum of the squares between the clusters multiplied by the weights of the features, as shown in Equation 1. The clusters are determined by maximizing this sum.

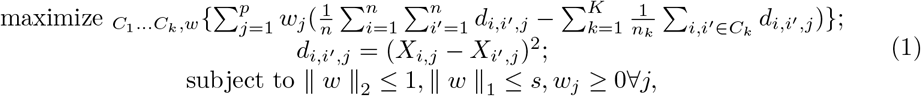

where *X* is a matrix n (sample) ×*p* (feature), *i* is the number of samples, *j* is the number of features, *w* is the weight of the feature, *w_j_* is the weight of the *j*-th feature, *k* is the number of clusters, *n_k_* is the number of samples in the *k*-th cluster number, and *C_k_* is the *k*-th cluster. The parameter *s* is the sum of the weights of the features. If *s* is large, more features are selected, and if *s* is small, fewer features are selected.

Four regression algorithms were used, including d) Support Vector Regression (SVR), e) RF, f) PLS, and g) sparse PLS. d) SVR transforms the input space with a kernel function (e.g., linear, polynomial, radial basis function, sigmoidal, and Gaussian). In the deformed space, SVR minimizes the absolute value of weight coefficients and error simultaneously to obtain a regression equation to avoid overfitting. The epsilon tube is defined, such that if the absolute value of the error is less than ε, there is no penalty in the training loss function.

e) RF is an ensemble learning algorithm that builds many decision trees during training and adopts the most common solution. For regression problems, RF returns the average predictions of the individual decision trees. RF is an extension of the bagging method. While decision trees consider all features, RF ensures a low correlation between decision trees by randomly selecting a subset of samples and features. Thus, it avoids the overfitting problem of decision trees and builds a discriminative method with high generalizability. RF has two primary hyperparameters. These include the number of trees to build and the maximum number of features considered at each leaf in the tree.

f) PLS uses a statistical method called principal component regression. While principal component analysis (PCA) extracts the principal components so that the explanatory features are maximized, PLS extracts the principal components so that the covariance between the explanatory and objective features is maximized. It can be considered a supervised version of PCA. PLS is gaining popularity in metabolomics and other *omics* analyses because it is particularly suitable for when the number of predictor features is greater than the observed ones and there is multicollinearity among the training data. g) Chun and Keles [8] proposed sparse PLS to address the complexity of the interpretability of high dimensional data where conventional PLS is commonly used. Sparse PLS simultaneously discriminates the given data and reduces the discriminant features by imposing sparsity using *L*_1_ penalty on the direction vector. The generalized regression equation used in sparse PLS, defined in Equation 2, repeats the solves *w* and also solves c with fixed *w*, in which c is a surrogate of the original direction vector *w* [8].

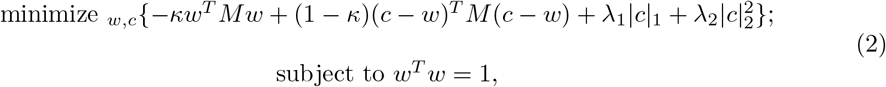

where, *M* = X^Y^*YY^T^* (X: the predictor matrix, *Y*: the response matrix). Small *κ* can reduce the local minimum risk. *L*_1_ penalty facilitates the sparsity, and *L*_2_ penalty deals with the potential singularity of *M* when solving for *c*.

### 2.2 Data selection

The MTBLS92 dataset [24] from MetaboLights [15] was used in this study. This dataset was obtained from a multicenter, randomized, phase III trial in which patients with breast cancer were randomly assigned to one of the following neoadjuvant chemotherapies (NACs).

4×epirubicine+cyclophosphamide (EC) → docetaxel (D)
4×EC → 4×D/capecitabine (C)
4×EC → 4×D→ C

Serum lipidomic samples were collected before NAC (baseline, [BL]) in the non-fasting state after NAC at the time of surgery. Serum lipid levels were quantified by liquid chromatography-mass spectrometry (LC-MS) and fatty acids by gas chromatography (GC) and a flame ionization detector (FID) detector. Substances were quantified based on the peak intensity ratio using internal standards spiked into each sample and identified using an internally-developed library search. The dataset consisted of 253 individuals and 143 metabolites. Unidentified metabolites were omitted from the analysis beforehand. We refer the reader to the original paper for details of the study design, sample collection, and processing [24]. The number of samples used in this study is listed in Table 1.

**Table 1.**
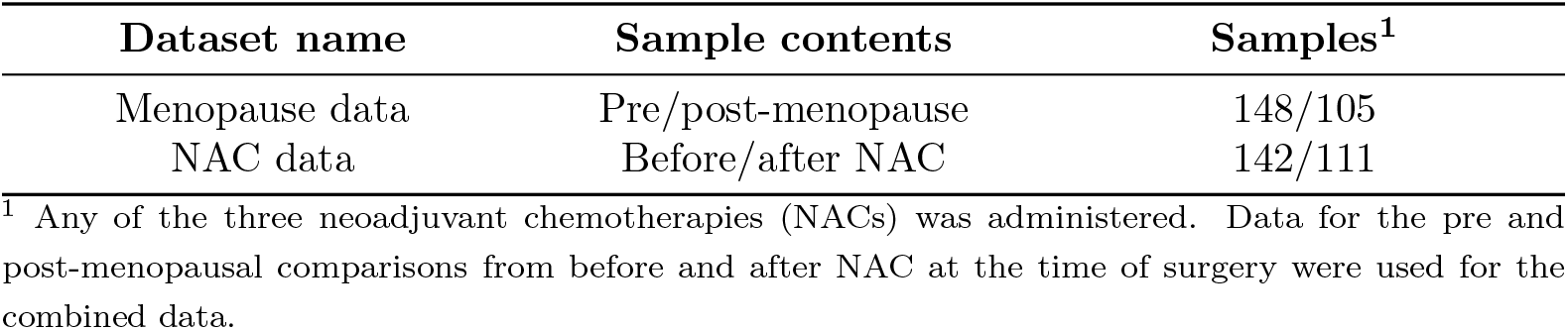
Number of samples

| Dataset name | Sample contents | Samples<sup>1</sup> |
| --- | --- | --- |
| Menopause data | Pre/post-menopause | 148/105 |
| NAC data | Before/after NAC | 142/111 |

<sup>1</sup> Any of the three neoadjuvant chemotherapies (NACs) was administered. Data for the pre and post-menopausal comparisons from before and after NAC at the time of surgery were used for the combined data.

### 2.3 Evaluation indicators

To compare the performance of the algorithm, we calculated three basic evaluation measures from the confusion matrix: accuracy, sensitivity, and specificity. In each experiment, post-menopause and after NAC cases were used as positive cases. The true positive (TP) and false negative (FN) ratios were those classified as positive and negative among positive cases. The true negative (TN) and false positive (FP) were the ratios classified as negative and positive among the negative cases. Accuracy was calculated by (TP + TN) / (TP + TN + FP + FN), sensitivity by TP / (TP + FN), and specificity by TN / (TN + FP). Pairwise t-tests with Bonferroni correction were used to evaluate the difference in quantitative values between positive and negative cases.

Because the output values of the regression algorithm are continuous scores, the classification accuracy depends on a single defined threshold. Therefore, a receiver operating characteristic (ROC) curve was plotted over the entire threshold for a more general evaluation, showing the trade-off between sensitivity and specificity. The area under the curve (AUC) was calculated with a 95% confidence interval (CI). The AUC can be interpreted as the probability of ranking a random positive instance higher than a random negative instance. Mann-Whitney U-test was performed for AUC (*P* < 0.05). This study also used the ROC curve to determine the optimal cut-off point to convert continuous scores to binary ones, which is described in detail in section 2.5.4.

Computational costs are essential when evaluating the practical applications of algorithms. Thus, we calculated the runtime required to train each algorithm. All calculations were performed on a MacBook Air, 1.1 GHz Intel Core i5.

### 2.4 Extraction of metabolites that contributed to the discrimination

Sparse K-means and each regression algorithm scored metabolites according to the contribution to the discrimination. The feature importance score for the sparse K-means is the weight of each feature. The sum of feature weights was determined by hyperparameter *s* in Equation 1. In linear SVR, features used for training are assigned weights proportional to their predictive capabilities. For RF, the Gini coefficient was calculated for each feature based on the Gini impurity [4]. In other words, feature importance was calculated based on the degree to which Gini impurity was reduced by splitting by a certain feature when constructing the decision tree. In PLS, the VIP score measures the importance of each feature in the projection, calculated as a weighted sum of the squared correlations between the PLS components and the original features. In sparse PLS, regression coefficients were used.

Each feature was ranked to compare the selected metabolites that highly contributed to the discriminations among the five algorithms. First, each feature was assigned a rank according to the importance score. Features with the same importance score were assigned the average rank. For each algorithm, the average feature rank was calculated by repeating 10-fold cross-validations (CV) with different randomizations 100 times. Finally, the features ranked in the top 50 on average across all trials were compared between algorithms.

### 2.5 Preprocessing and classification process

#### 2.5.1 Data splitting and preprocessing

Hilvo et al. investigated a difference in the metabolite profiles of patients who received NAC before and after treatment. They also analyzed the difference between samples from pre/post-menopausal patients [24]. This study used two classifications: (1) pre/post-menopause and (2) before and after NAC.

The performance of each algorithm was evaluated via a stratified 10-fold CV, which was repeated 100 times with different randomization in each repetition. The stratified splitting in the CV ensured that the class distribution of each fold was approximately the same across folds. The same data was used for all algorithms for a fair comparison. However, in regression algorithms, a nested stratified 10-fold CV approach was used to select classification thresholds.

Only one data value was missing: the triglycerides (50:5) data for one patient. It was complemented using the min/2 of each feature. Values from the training data complemented missing values in the test data. Furthermore, the values for each sample were subjected to a log2-transformation and Z-score normalization, followed by Z-score normalization for each feature. For the Z-score normalization of the test data, the mean and standard deviation (std) of the training data were used.

#### 2.5.2 Feature selection using the filter method and sparse modeling

Filtering and feature selection using sparse modeling were compared. Filtering is frequently used in metabolomic data analyses. Univariant filtering evaluates each feature individually, and the top *N* features (N is predefined) are utilized for subsequent analyses. Here, two filtering methods were employed: (1) filtering based on the variance of each metabolite, i.e., unsupervised, and (2) filtering based on the AUC values of each feature, i.e., supervised. These feature selections were conducted before the clustering and regression.

Meanwhile, sparse modeling simultaneously performs feature selection and algorithm training. We compared the classification accuracy of these three feature selection methods under the condition where half of the metabolites were selected. Because the location of the true biomarkers is unknown, classification accuracy was used as an indicator to evaluate the quality of the selected feature subset. For the sparse K-means and sparse PLS, the hyperparameter controls the number of features to be selected. Hyperparameter *s* ranged between 1.5 and 10 with 20 divisions for sparse K-means, and hyperparameter *eta* ranged between 0.1 and 0.9 with 20 divisions for sparse PLS. The hyperparameters were set to select approximately half of the metabolites in each sparse modeling (i.e., *s* = 7.0 and *eta* = 0.5). For AUC filtering, the procedure described in section 2.5.1 was used. For variance-based filtering, the metabolite concentrations were converted using log2-transformation and Z-score normalization.

#### 2.5.3 Clustering process

We used K-means and HC, two of the most popular clustering algorithms. Sparse K-means, proposed by Witten and Tibshirani [55], was also used to evaluate the effectiveness of sparse modeling compared to conventional non-sparse modeling. We labeled the two clusters obtained from the training data in such a way that the accuracy of the training labels was increased. The test data were predicted to have the classification label with the closest Euclidean distance to the cluster center of gravity. Sparse K-means simultaneously performs feature subset optimization with training. Therefore, only features selected during training were used for the computation of the center-of-gravity vector and testing. Hyperparameter *s* defining sparsity was determined according to the method described in section 2.5.2. For HC, we used agglomerative hierarchical clustering, which is computationally inexpensive, and the Ward method to measure the distance between clusters.

#### 2.5.4 Regression process

The three most popular regression algorithms, SVR, RF, and PLS, were evaluated and compared with sparse PLS, proposed by Chun and Keles [8]. The classification threshold for each test was selected via a nested stratified 10-fold CV in the inner loop to classify the binary class. After calculating continuous values for each regression analysis using the training data, the ROC curves were depicted using the validation data. The furthest vertical distance from the diagonal on the ROC curve (also called the Youden index [59]) was used as the best threshold. The average value of the Youden index calculated from 10-inner loops was used as the classification threshold to compare classification accuracy. The hyperparameters for each regression algorithm used the default values provided in the package, which produce the baseline performance. For liner SVR, an insensitivity coefficient *ε* = 0.1 was set to control the region where errors are ignored, and a regularization coefficient *C* =1.0 was set to control the strength of regularization and prevent overlearning. For RF, the number of trees was set to 100, and the maximum number of features was considered at each leaf in the tree. For PLS and sparse PLS, the number of components to keep was set to 2. Hyperparameter eta defining sparsity was determined according to the method described in section 2.5.2.

### 2.6 Software implementation

In our experiments, we used a MacBook Air 1.1 GHz Intel Core i5 16 GB RAM with Python 3.9.1 [53] and R 4.2.0 [51]. The K-means method was computed using the standard R package “stats” (version 4.0.3) [51]. The sparse K-means method was computed using the R package “sparcl” (version 1.0.4) [56], and sparse PLS was computed using R package “spls” (version 2.2-3) [9]. Other algorithms were calculated using the Python package “scikit-learn” (version 1.0.2) [44]. For multiple tests, Python package “scikit-posthocs” (version 0.7.0) was used [52]. The program used for the classification problems in this study will be made available upon request.

## 3 Results

Here, we evaluated the commonly used algorithms and sparse modeling in metabolomic data analysis. The comparison of classification accuracies and computational costs are presented in section 3.1. The comparison of selected features are presented in section 3.2. The sparse modeling methods were repeatedly conducted to analyze the stability of the feature selection, and the comparison of selected features with those from filtering methods are presented in section 3.3.

### 3.1 Classification performance comparison

The classification performance of sparse and non-sparse modeling was compared using three basic metrics calculated from the confusion matrix: accuracy, sensitivity, and specificity. Figure 2 shows the mean and 95% confidence interval (CI) of the metrics for each algorithm used in this study. Post hoc pairwise comparisons with Bonferroni correction were used to compare the performance of each algorithm. Sparse modeling (i.e., sparse K-means and sparse PLS) showed similar accuracies as another non-sparse modeling in menopause classification. Furthermore, sparse PLS showed significant differences (*P* < 10^-100^) from K-means and sparse K-means in NAC classification.

**Figure 2.**
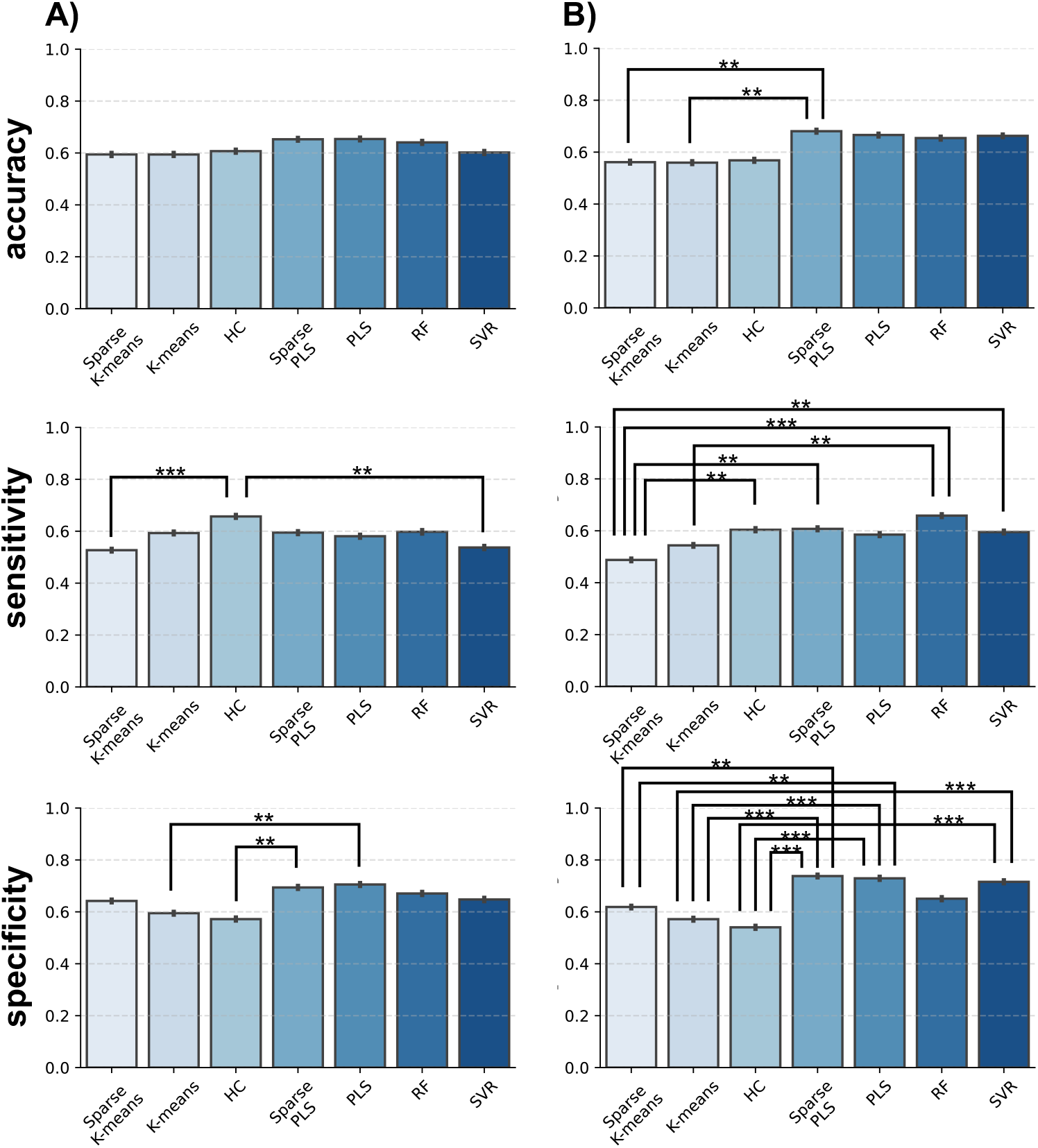
Comparison of accuracy-based classification performance metrics: (A) pre/post-menopause classification results; (B) before and after NAC classification results. Accuracy indicates the number of correctly classified samples. The sensitivity indicates the power to discriminate post-menopause and after NAC cases. Specificity indicates the ability to detect menopause or after NAC cases. Error bars indicate the 95% CI obtained via 10-fold CV, repeated 100 times with different randomization in each repetition. All data were analyzed using pairwise t-tests with a Bonferroni correction to compare the different algorithm performances (**: *P* < 10^-100^, ***: *P* < 10^-150^)

The ROC curve was used for the regression algorithms to evaluate the balance between specificity and sensitivity obtained across some thresholds (Figure 3). In the menopause classification problem (Figure 3a), the AUC for sparse PLS was 0.7177 (95% CI, [0.7112, 0.7240]), 0.7162 for PLS (95% CI, [0.7097, 0.7223]), 0.7024 for RF (95% CI, [0.6959, 0.7087]), and 0.6440 for SVR (95% CI, [0.6368, 0.6505]). In the NAC classification problem (Figure 3b), the AUC for sparse PLS was 0.7315 (95% CI, [0.7253, 0.7380]), 0.7080 for PLS (95% CI, [0.7016, 0.7147]), 0.7206 (95% CI, [0.7146, 0.7266]), and 0.7220 for SVR (95% CI, [0.7157, 0.7285]). For both classification problems, sparse PLS had the best mean AUC. All algorithms showed significantly higher ROC values (*P* < 0.05, Mann-Whitney U-test) than random predictions.

**Figure 3.**
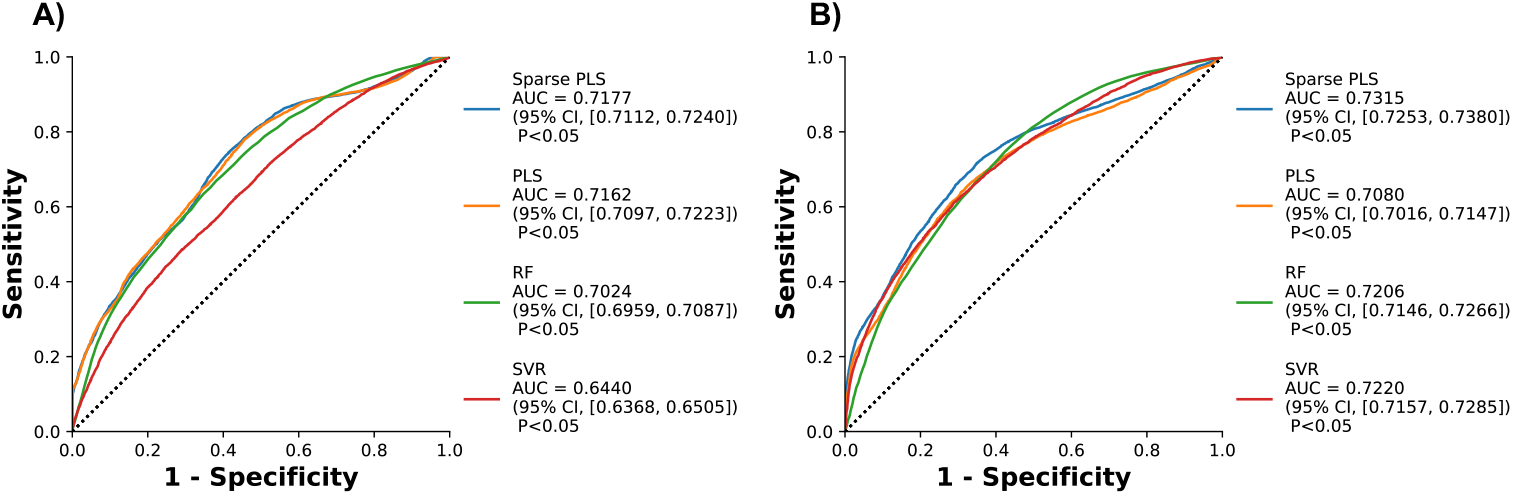
ROC curves showing sensitivity and specificity of the classification results. (A) Classification results for pre and post-menopause. (B) Classification results for before and after NAC. 95% CIs were obtained via 10-fold CV, repeated 100 times with different randomization in each repetition. The dotted diagonal lines represent a completely random prediction. The Mann-Whitney U-test was used to compare prediction results with the random predictions.

The numerical experiments measured the runtime as a computational cost (Figure 4). HC and PLS showed the shortest runtime and less variability in multiple trials than the other algorithms. Sparse modeling (i.e., sparse K-means and sparse PLS) was shown as computationally expensive compared to the traditional standard algorithms (i.e., K-means and PLS). However, the increase in runtime for sparse PLS relative to PLS was less than that for sparse K-means relative to K-means. Moreover, sparse PLS was found to be computationally less expensive than RF and SVR, even though feature selection is performed simultaneously with algorithm training.

**Figure 4.**
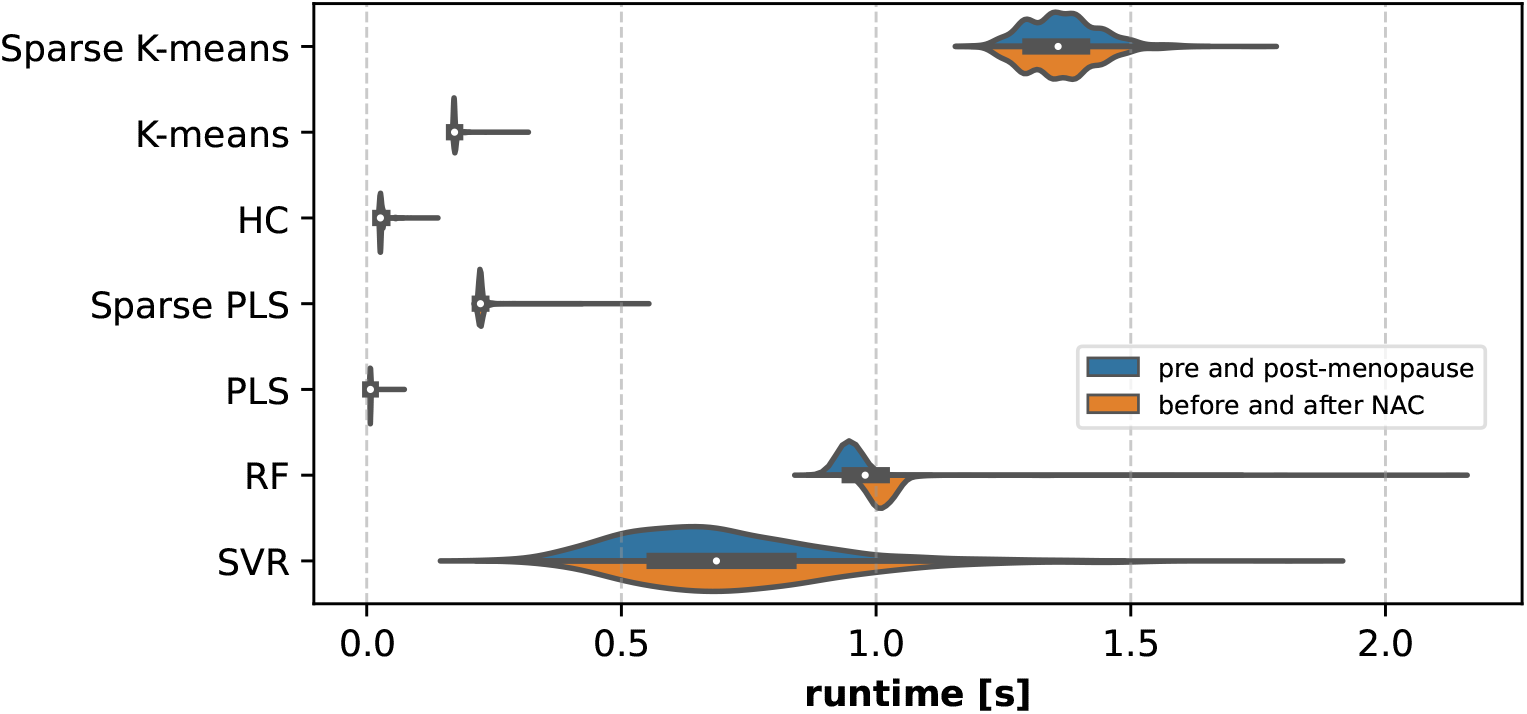
Comparison of runtime among three clustering and four regression algorithms. The algorithm training time was measured as the runtime for each trial. The dataset is split via a 10-fold CV, repeated 100 times with different randomization in each repetition. The thickness of the region indicates the distribution of the number of trials. The distributions shown in blue are the results of pre/post-menopause classification and the distributions shown in orange are before and after NAC classification. The black box-and-whisker diagram in the center shows the runtime distribution for the two combined classification problems, and the white dot indicates the median value.

### 3.2 Identification of metabolites that contributed to the discrimination

Sparse PLS and four regression algorithms were used to score metabolites that contributed to the discrimination during training. All 143 metabolites ranked according to feature importance score corresponding to each algorithm were obtained and used to identify the frequently selected metabolites. For each algorithm, the average rank of each metabolite was calculated across all trials. The top 50 metabolites were extracted as features that highly contributed to discrimination. The results are shown in Figure 5.

**Figure 5.**
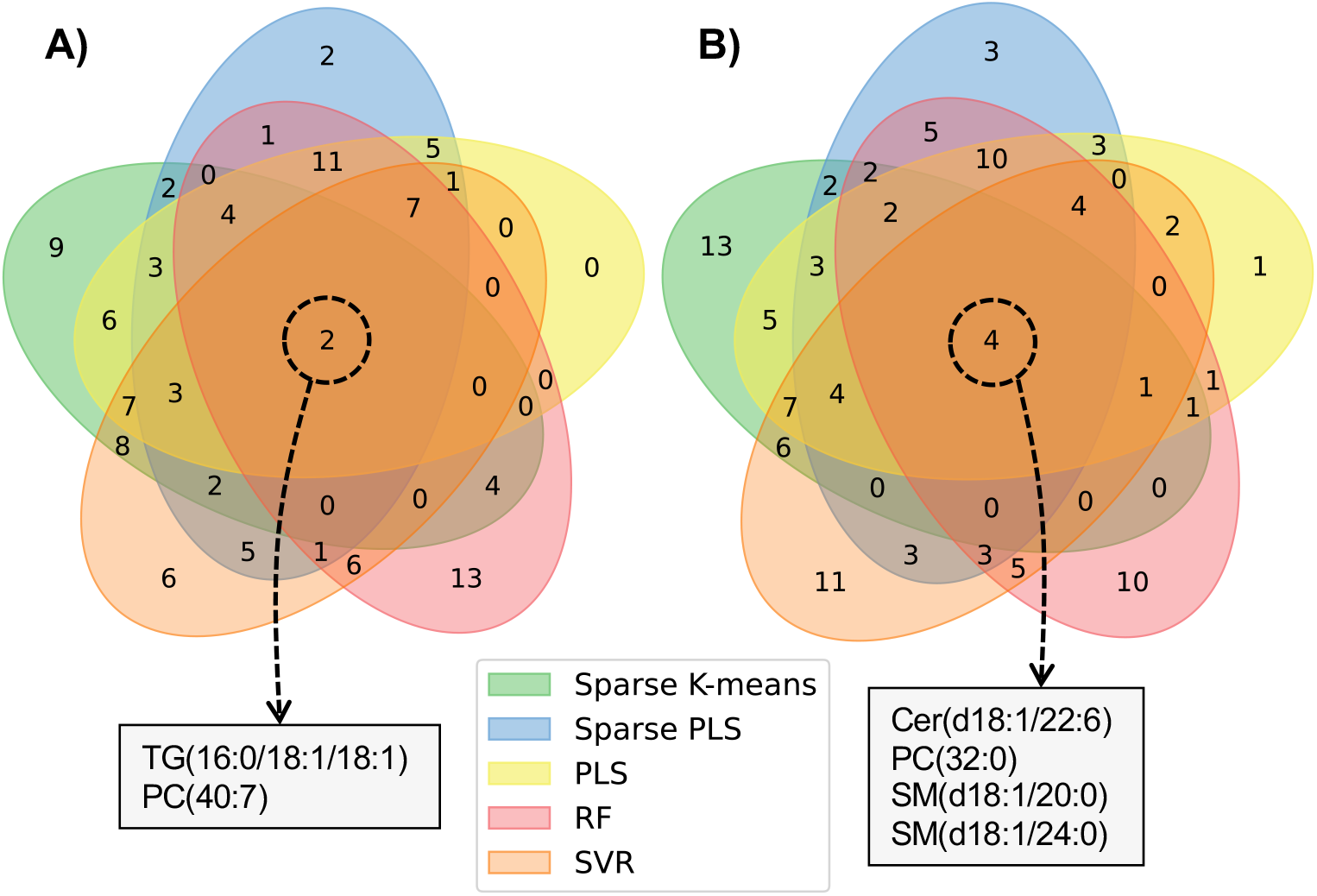
Venn diagram showing the identified metabolites that contributed to discrimination in the five algorithms. (A) Discrimination results of the menopause cases. (B) Discrimination results of NAC cases. Each metabolite is scored based on the feature importance obtained during training. Metabolites ranked within the top 50 on average across all trials are counted as metabolites that highly contributed to discrimination. The center of the Venn diagram shows the number and detail of selected metabolites common to all five algorithms.

For menopause discrimination, two metabolites, triacylglycerol (TG) (16:0/18:1/18:1) and phosphatidylcholine (PC) (40:7) were commonly selected in all five algorithms. For NAC discrimination, four metabolites, ceramide (Cer) (d18:1/22:6), PC(32:0), sphingomyelin (SM) (d18:1/20:0) and SM(d18:1/24:0), were commonly selected in all five algorithms.

Furthermore, we analyzed the inter-algorithm similarity of the metabolite sets that contributed to the discrimination. We counted the number of top 50 metabolites common to the two algorithms (Figure 6). For both discrimination problems, the two sparse modeling algorithms (i.e., sparse K-means and sparse PLS) shared more than half of the metabolites with PLS. For sparse PLS, more than half of the metabolites were also shared with RF.

**Figure 6.**
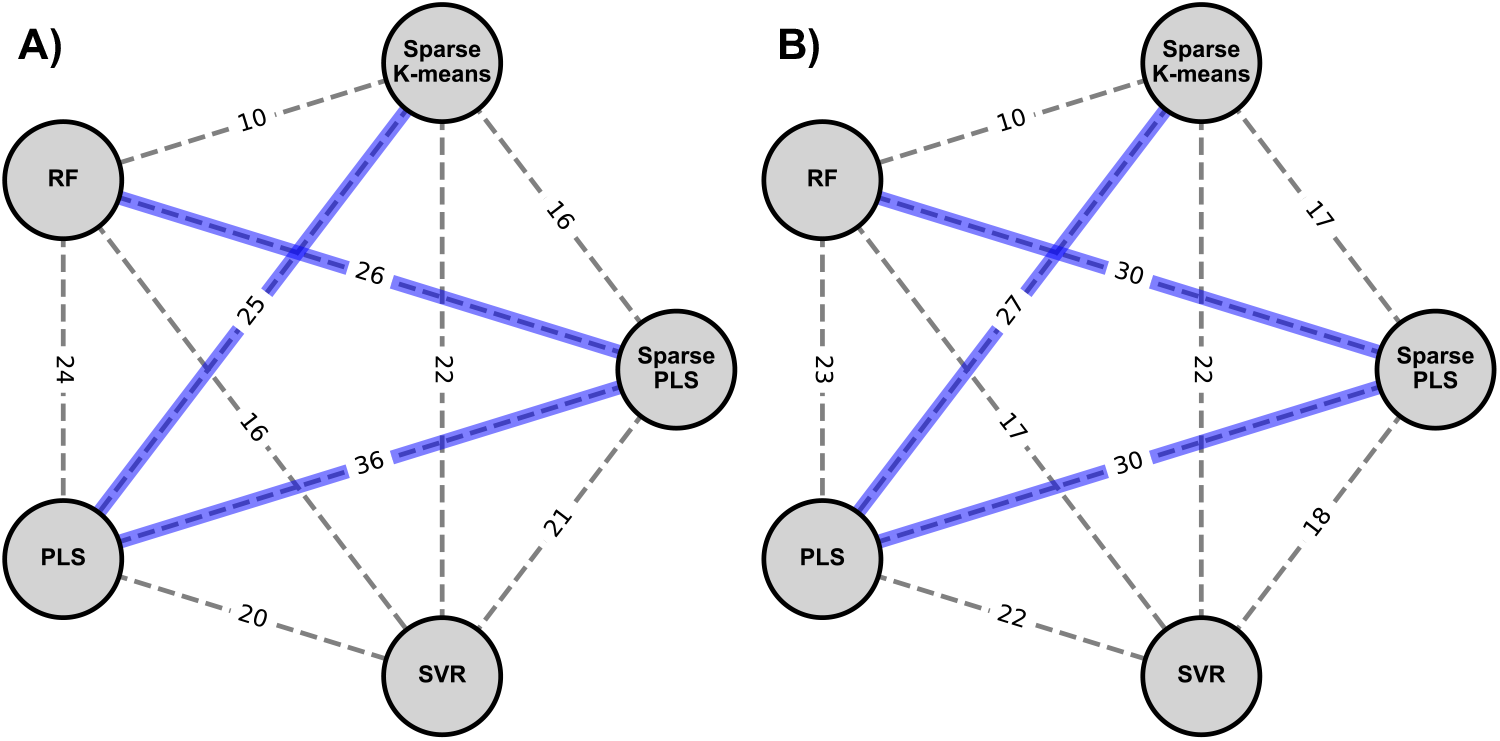
Weighted network graphs showing the number of metabolites commonly contributing to discrimination for each algorithm pair. (A) Discrimination results of the menopause cases. (B) Discrimination results of NAC cases. Each metabolite is scored based on the feature importance score obtained during training. Metabolites ranked within the top 50 on average across all trials are extracted as metabolites contributing to discrimination. The number on the edges represents the number of metabolites common to the top 50 in the two algorithms. Furthermore, blue edges represent pairwise comparisons where more than half are the same metabolite.

### 3.3 Stability of metabolite selection by sparse modeling and its effectiveness

Sparse modeling enables both feature selection and model training. Here, the stability of the feature selection of sparse modeling was evaluated. The relationship between the number of selected features and the hyperparameter that controls the sparsity was evaluated using the NAC dataset (Figure 7). In sparse K-means, the number of selected metabolites increased along with the increase of *s*, from an average of 4 metabolites at *s* = 1.5 to an average of 143 metabolites at *s* = 10.0. In sparse PLS, the number of selected metabolites increased as eta decreased, from an average of 5 metabolites at *eta* = 0.9 to an average of 142 metabolites at *eta* = 0.1. Sparse K-means showed the largest std at *s* = 9.55 (std = 3.79), and sparse PLS showed the largest std at *eta* = 0.48 (std = 5.54). This variation was higher at approximately *eta* = 0.48 and lower when eta was far from 0.48. Overall, Sparse K-means showed smaller variations, which indicated a more stable variation in the number of selected metabolites relative to hyperparameters compared to sparse PLS.

**Figure 7.**
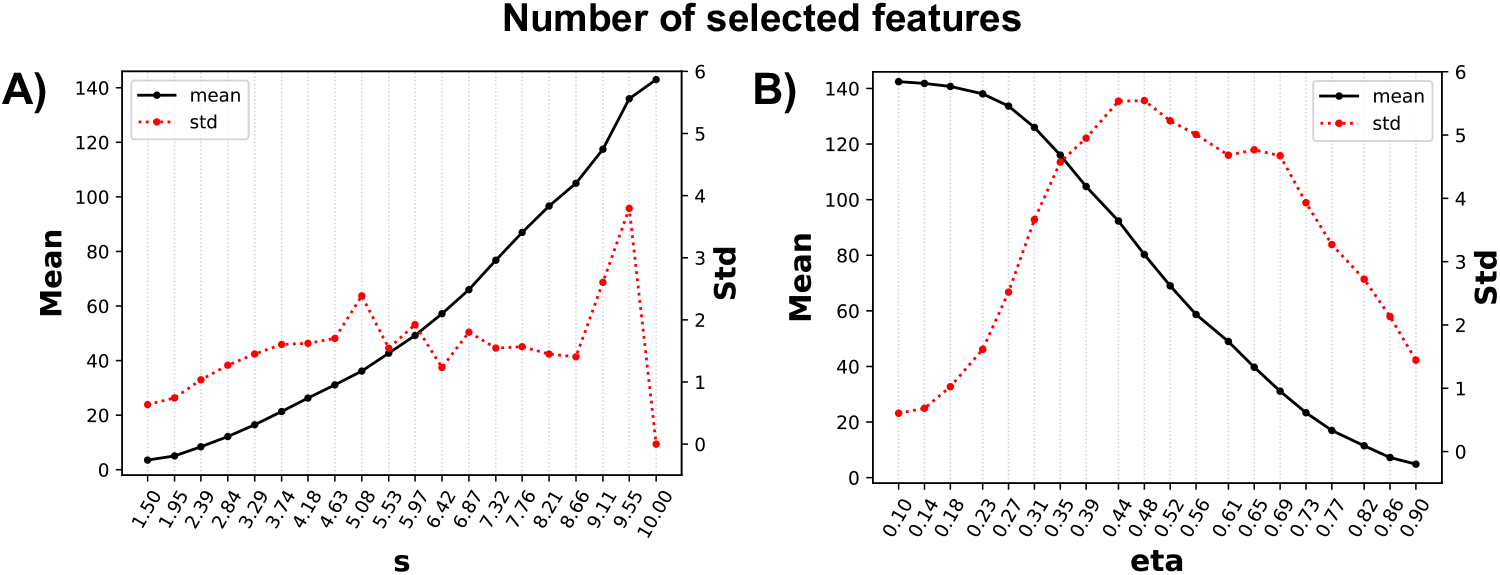
The mean and standard deviation (std) of selected metabolites for hyperparameters that define sparsity in two sparse modeling methods. The number of selected metabolites for the (A) sparse K-means and (B) sparse PLS. In sparse K-means, hyperparameter *s* ranged from 1.5 to 10, with 20 divisions. In sparse PLS, hyperparameter eta ranged from 0.1 to 0.9, with 20 divisions. The NAC dataset was split via a 10-fold cross-validation, repeated 100 times with different randomization in each repetition. The solid black line represents the mean number of the selected metabolites, and the dotted red line shows the std of the selected metabolites.

The feature selection of sparse modeling was compared to feature filtering. Variance and AUC were used for univariant feature filtering, commonly used in metabolomics, as described in section 2.5.2. The comparison of classification accuracy using NAC data is depicted in Figure 8. The sensitivity of K-means with AUC-based filtering was significantly higher (*P* < 10^-100^) than K-means with variance-based filtering and sparse K-means. The accuracies were not significantly different between the embedded method (i.e., sparse modeling) and the filtering method by AUC and by variance, indicating that sparse modeling showed similar classification performance compared to the filtering methods.

**Figure 8.**
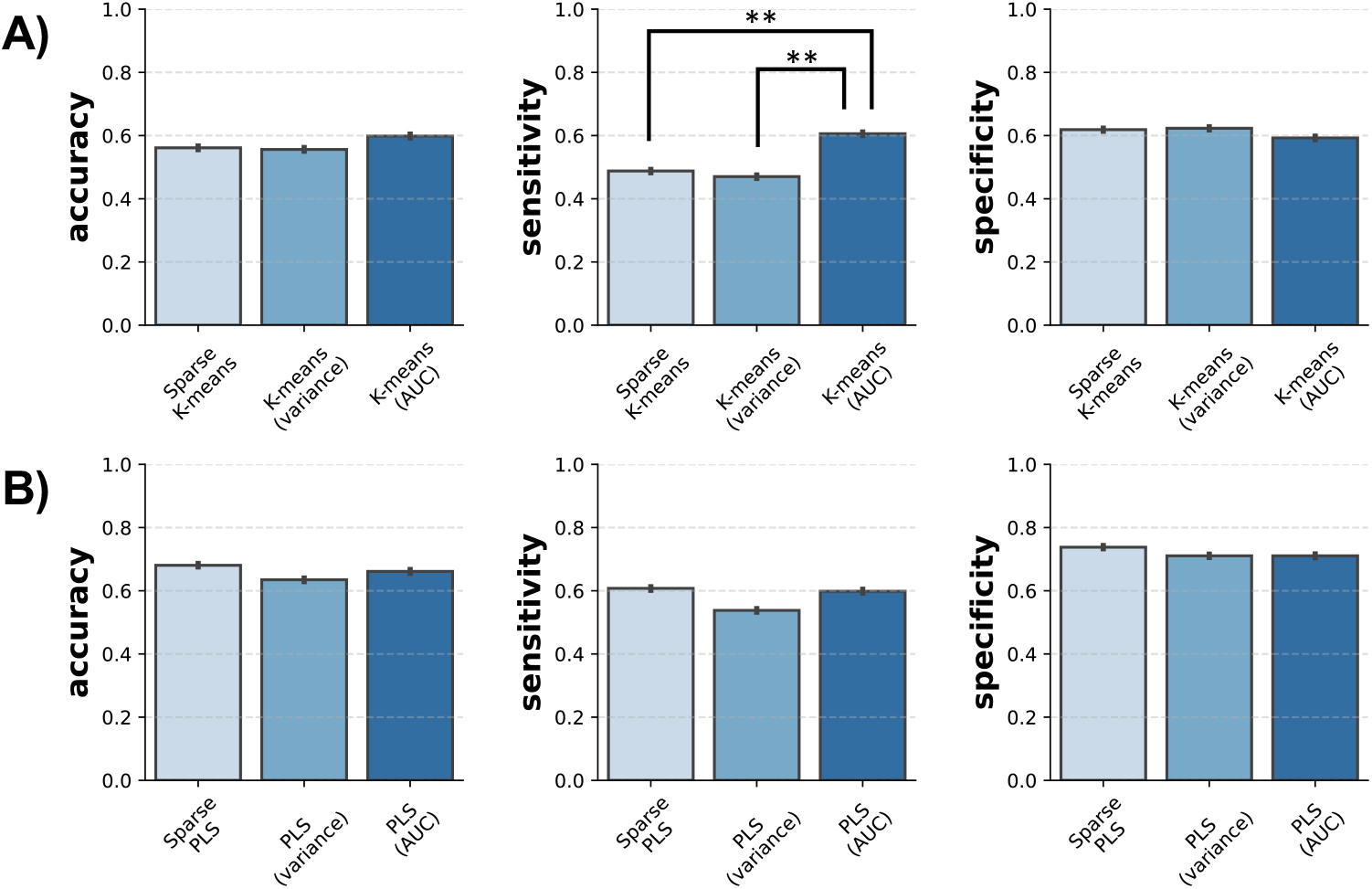
Comparison of classification performance of feature selection using sparse modeling and pre-screening feature filters. Results for the (A) sparse K-means and (B) sparse PLS. Error bars indicate the 95% CI obtained via 10-fold cross-validation, repeated 100 times with different randomization in each repetition. NAC classification data were used. All data are analyzed using pairwise t-tests with a Bonferroni correction to identify algorithm performances that are different from each other (**: *P* < 10^-100^).

## 4 Discussion

This study compared sparse modeling methods, including sparse K-means and sparse PLS, with other commonly used analytical algorithms in metabolomics research. The numerical experiment was conducted using two comparisons (menopause and NAC) of serum lipidomics profiling data of patients with breast cancer. The classification accuracies and computational costs were compared. The sparse modeling algorithms that used a subset of features showed equal to or better accuracies than other algorithms that used all features. Among the five algorithms, two metabolites in the menopause classification and four metabolites in the NAC classification were identified as the common metabolites with a high contribution to discrimination. The selected features by sparse modeling were consistent with the VIP score of PLS as the current gold standard feature selection tool in the metabolomics field. The repeated trials revealed the stability of the feature selection using sparse modeling, and feature selection by sparse modeling resulted in similar quality feature sets with the filtering method.

Even though sparse modeling used a subset of features, its classification performance was equal to or better than non-sparse modeling using all features. This suggests that sparse modeling may be a useful classification algorithm in metabolomics analyses. However, sparse modeling (i.e., sparse K-means and sparse PLS) require more computational costs for algorithm training than standard non-sparse modeling (i.e., K-means and PLS). This is because sparse modeling involves the simultaneous optimization of clustering and regression equations and the optimization of features subsets. Sparse PLS achieves sparsity by adding *L*1 penalty to the objective functions of the direction vector. Thus, its computational cost is greater than that of PLS but still lower than the other non-sparse modeling algorithms (i.e., SVR and RF).

A typical metabolomics study identifies biomarker sets that correlate with the phenotype investigated. We explored the metabolites that highly contributed to the discrimination of each algorithm. The algorithms commonly selected PC(40:7) and TG(16:0/18:1/18:1) in the menopause classification, and Cer(d18:1/22:6), PC(32:0), SM(d18:1/20:0), and SM(d18:1/24:0) in the NAC classification as discriminant features. A plasma lipidomics study reported an elevated concentration of PC(40:7) in patients with breast cancer [7]. The disturbance in the lipid profile of plasma samples, including PC, was investigated, according to the menopausal status among healthy women [43]. TG(16:0/18:1/18:1) was one of triglycerides identified in the study. While not exactly the same as this metabolite, the change in TG metabolites depends on the menopausal status [43]. Cer(d18:1/22:6) is a ceramide metabolite that is a combined form of sphingosine and fatty acids, and is usually found in high concentrations in the cell membrane. Ceramide promotes cell apoptosis or senescence by modulating the activity of oncogenes [16]. The elevated concentration of these metabolites in breast cancer tissue, especially estrogen-positive tumor tissue, has been reported [46]. The lipid profiles vary among breast cancer subtypes [14]. PC(32:0) levels were elevated in HER2-positive breast cancer tissue compared to the HER2-negative tissues [31]. This metabolite was also quantified in triple-negative breast cancer tissue; however, no correlation with the expression level of Ki67, a proliferation marker of breast cancer, was observed [25]. Sphingomyelin, such as SM(d18:1/20:0) and SM(d18:1/24:0), in serum samples between the patients of invasive ductal carcinoma type of breast cancer and matched healthy controls were different [10]. Thus, the metabolites commonly selected by various methods are known for their relationships with menopausal status and subtypes of breast cancer, which may reflect the efficacy of the chemotherapies.

In breast cancer patients treated with NAC, the serum lipidomic profile remains unchanged, but changes in thiobarbituric acid reactive substances (TBARS) and super-oxide dismutase (SOD) indicate that oxidative stress in the body is significantly altered [20], resulting in a change in TG levels. Epirubicin, an anthracycline anticancer drug, was included in the NAC. It has been reported that some SMs may contribute to the activation efficiency of anthracycline uptake [54]. PC has been shown to protect against peripheral neurotoxicity, one of the side effects caused by docetaxel [32]. Changes in the serum lipidomic profile with NAC treatment are also speculated. In breast cancer subtypes, triple negative tumors contain more specific PCs [25], suggesting that the baseline may differ in each subtype rather than uniformly changing in all cases. In contrast, there are reports that serum PC levels in breast cancer patients decreases during anticancer drug treatment [36]. PC can be used as a marker for predicting cervical cancer susceptibility to NAC [58]. Therefore, changes in the serum lipidomic profile may vary from individual to individual.

We focused on the number of common metabolites in the selected feature subsets between the two algorithms. We found that sparse K-means and sparse PLS overlap with the majority of features chosen by the VIP of PLS. The selection results of sparse PLS were also similar to those of RF. In metabolomics, PLS is frequently used for discrimination and dimensionality reduction [3, 42]. Using a semi-artificial database with known biomarker locations, Manon and Govaerts [41] showed that the VIP score of PLS is often the best method for discrimination. RF is reported to be insensitive to noise and has little or no overfitting. Therefore, RF often shows a more suitable classification and biomarker selection performance than the widely preferred PLS and SVM [6, 47]. RF has been used as a biomarker selection tool for breast cancer [26], hepatocellular carcinoma [19], and metabolic syndrome [38]. The similarity of the feature selection by the two sparse modeling methods to that of PLS and RF suggests that sparse modeling is helpful for biomarker identification in metabolomics studies.

One concern in using sparse modeling for feature selection is the possibility that the size of the selected feature subset may vary with iterations of the algorithm. Therefore, the stability of feature selection was evaluated by globally perturbing the hyperparameters that define sparsity and by iterating training 1,000 times at each hyperparameter setting. A comparison of the two sparse modeling methods revealed that sparse K-means showed a lower variability in the number of features selected than did sparse PLS. Sparse PLS showed a greater variation in the number of features chosen as *eta* = 0.48 was approached. On the other hand, the variation in the number of features selected using sparse K-means was robust to hyperparameters. At *s* = 9.55, variation increased to std = 3.79, but this effect can be ignored in many contexts where feature selection is performed because there are few situations to omit a few features.

In addition, to evaluate the quality of the selected feature subset, feature selection and two unsupervised and supervised filtering (i.e., filtering by variance and by AUC) were compared using sparse modeling. The feature subset selected by sparse modeling showed similar classification accuracy to those selected by filtering. However, K-means with pre-screening of features by AUC filtering showed better sensitivity than sparse K-means. This result may be explained by the fact that sparse K-means is essentially an unsupervised learning algorithm and was pre-trained with supervised filtering by AUC. Based on these results, feature selection by sparse modeling is a valuable alternative to conventional filtering methods.

Our research has several limitations. Our results were evaluated using only two classifications and do not include a comparison with artificial neural networks, a popular machine learning algorithm. In sparse modeling, objective hyperparameter adjustment methods have been proposed to determine feature subset size. Originally, in sparse K-means, it is necessary to measure the gap statistic to determine the hyperparameter *s* that defines the sparsity [18, 37, 55].

## 5 Conclusion

In this study, the performance of basic clustering and regression algorithms and their sparse models for metabolomics data analysis were compared. The results show that sparse modeling performs as well as or better than conventional algorithms, even though it uses only a subset of features. The second major finding was that both regression and clustering algorithms of sparse modeling focused on a similar subset of the current benchmark, the VIP-based feature selection in PLS. Therefore, the results of this study indicate that sparse modeling is useful for biomarker identification because it is able to identify the smallest set of strongest predictors associated with a particular condition. These findings contribute to the rapidly expanding field of sparse strategy in *omics* analysis.

